# Mirdametinib and abemaciclib cooperate in atypical teratoid rhabdoid tumor to decrease proliferation and suppress tumor growth

**DOI:** 10.64898/2026.03.24.714018

**Authors:** Jinming Liang, Yiming Deng, Anupa Geethadevi, Kristen Malebranche, Tyler Findlay, Charles Eberhart, Jeffrey Rubens, Eric Raabe

**Affiliations:** Department of Biochemistry and Molecular Biology, Johns Hopkins University Bloomberg School of Public Health, Baltimore, MD; Division of Pediatric Oncology, Department of Oncology, Johns Hopkins University School of Medicine, Baltimore, MD; Division of Cell Biology, Johns Hopkins University School of Medicine, Baltimore, MD; Division of Neuropathology, Department of Pathology, Johns Hopkins University School of Medicine, Baltimore, MD

**Keywords:** MEK inhibitor, cyclin dependent kinase, ERK, pediatric brain tumor

## Abstract

Atypical teratoid rhabdoid tumor (ATRT) is a malignant brain tumor of children that has an overall survival of less than 40 percent even with aggressive therapy. We identified upregulation of the mitogen activated protein (MAP) kinase pathway in ATRT. The novel, brain-penetrant MEK inhibitor mirdametinib inhibited the growth of ATRT cell lines in culture at nanomolar concentrations. Mirdametinib suppressed proliferation as measured by BrdU incorporation and induced apoptosis as measured by cPARP and Annexin V staining. Monotherapy with mirdametinib extended the life of mice bearing orthotopic xenografts. Combination therapy with the brain-penetrant cyclin dependent kinase 4/6 inhibitor abemaciclib further suppressed growth and BrdU incorporation in ATRT cell lines representing all molecular subgroups. Mirdametinib and abemaciclib combined to extend survival of mice bearing orthotopic ATRT xenografts. In conclusion, mirdametinib has single agent activity against ATRT and combines with abemaciclib to decrease proliferation and extend survival in orthotopic xenograft models of ATRT.

## 1. Introduction

Atypical teratoid rhabdoid tumor (ATRT) is most common malignant brain tumor of infants. Overall survival is poor, with only 37% of patients achieving cure^1,2^.

Therapies include surgery, high dose chemotherapy with stem cell transplant, and in many cases radiation. Side effects of these therapies are often severe^1,2^. New treatments are urgently needed.

ATRT is an epigenetically dysregulated tumor, with the hallmark mutation being alteration of SMARCB1, a member of the mammalian SWI/SNF (mSWI/SNF) complex. The mSWI/SNF complex promotes differentiation, and when SMARCB1 function is lost, the ATRT cells fail to differentiate properly and are maintained in a stem cell state by overactivation of the Polycomb B repressive complex^3,4^.

We have identified the stem cell factors LIN28A and LIN28B as being important drivers of brain tumors^5,6^. These stem cell factors bind to and prevent the maturation of the *let-7* tumor suppressing microRNAs. *let-7* miRNAs in turn bind to and suppress the translation of mRNAs associated with cell growth, invasion, and stemness^7,8^. We demonstrated that suppression of LIN28A led to downregulation of KRAS in ATRT, and subsequently showed that ATRT primary tumors had increased activation of the mitogen-activated protein kinase (MAPK) pathway^5^. We found that the MEK inhibitors selumetinib and binimetinib suppressed ATRT proliferation, induced apoptosis, and decreased the growth of heterotopic ATRT tumors^5,9^. However, the relatively poor brain penetration of these MEK inhibitors prevented their application to ATRT growing in the brain^10^.

Mirdametinib is a next generation MEK inhibitor that has improved brain penetration compared to prior MEK inhibitors^11^. Mirdametinib is FDA approved for treatment of plexiform neurofibromas in adults and children with type 1 neurofibromatosis_12_. Mirdametinib is currently being evaluated in a phase 2 clinical trial for pediatric low-grade glioma (ClinicalTrials.gov Identifier: NCT04923126). We hypothesized that mirdametinib would penetrate the brains of mice, suppress the MAP kinase pathway, and improve survival in mice models of ATRT.

## 2. Materials and Methods

### 2.1. Cell culture and Reagents

The ATRT cell lines of CHLA-04-ATRT, CHLA-05-ATRT, and CHLA-06-ATRT were kindly provided by Anat Erdreich-Epstein^13^. The ATRT cell line, BT37, was developed from a serially passaged xenograft derived from a patient with ATRT14,15. BT12-ATRT and CHLA266-ATRT cell line were acquired from Children’s Oncology Group cell repository_16_. MAF737 and BT16 were a gift from Venkatraman Lab (University of Colorado, Aurora, CO, USA)^17^. CHLA05 and CHLA06 cells were cultured at 37 °C in a humidified 5% CO_2_ incubator in EF medium composed of 70% DMEM (Cat#11965092, ThermoFisher Scientific) and 30% Ham’s F-12 (Cat#11765054, ThermoFisher Scientific), supplemented with, L-glutamine (1%; Cat#25030081, ThermoFisher Scientific), B27 supplement without vitamin A (2%; Cat#12587010, Gibco), EGF (20 ng/mL; PeproTech, Cat#AF-100-15-1MG), FGF-2 (20 ng/mL; Cat#AF-100-18B-1MG, PeproTech), and heparin (5 µg/mL; Cat#H3149-100KU, Sigma). BT37, MAF737 and BT16 cells were grown at 37 °C in RPMI medium supplemented with fetal bovine serum (10%, Cat#A5670801, ThermoFisher Scientific), L-glutamine (1%; Cat#25030081, ThermoFisher Scientific) in a humid atmosphere with 5% CO2. All cell lines were validated by short tandom repeat (STR) testing and were routinely tested for mycoplasma.

### 2.2. Antibodies and Western immunoblotting

The following primary antibodies were used for immunofluorescence or western blot: anti-β-actin (1:1000, Cat# 4967L, Cell Signaling Technologies), anti-p44/42 MAPK (Erk1/2) (1:1000, Cat# 9102L, Cell Signaling Technologies), anti-Phospho-p44/42 MAPK (Erk1/2) (Thr202/Tyr204) (1:1000, Cat#4370S, Cell Signaling Technologies), anti-Cleaved PARP (Asp214) (1:1000, Cat# 5625S, Cell Signaling Technologies), anti-Rb (4H1) (1:1000, Cat# 9309S, Cell Signaling Technologies) and anti-phospho-Rb (Ser807/811) (1:1000, Cat# 8516S, Cell Signaling Technologies). The following secondary antibodies were used for western blot: goat anti-mouse IgG, HRP conjugate (1:5000, Cat# 91196S, Cell Signaling Technologies), goat anti-rabbit IgG, HRP conjugate (1:3500, Cat# 7074S, Cell Signaling Technologies). ATRT cells were collected after treatment mirdametinib alone and in combination with abemeciclib, and lysed using lysis buffer containing RIPA buffer, protease and phosphatase inhibitors.

Protein quantification was done using the Bradford method (Cat#5000006, Bio-Rad Laboratories Inc., Hercules, CA, USA). A total of 20 µg of protein isolated from cells or mouse tissues was resolved on 4–12% Bis-Tris gels (Cat# NW04122BOX, ThermoFisher Scientific, Waltham, MA, USA) by electrophoresis at 120 V. Proteins were then transferred to PVDF membranes (Cat#1620177, Bio-Rad Laboratories Inc., Hercules, CA, USA) by the Trans-Blot Turbo Transfer System (Bio-Rad Laboratories Inc., Hercules, CA, USA) at 25 V for 10 min.

Membranes were blocked in 5% BSA (Cat#03116956001, Sigma-Aldrich, NJ, USA) for 1 hour and incubated with primary antibodies overnight at 4°C. Then incubated with secondary antibodies for 1 h at room temperature. Then Blots were developed using enhanced chemiluminescence (ECL; Cat#NEL103001EA, Revvity Inc., Waltham, MA, USA; ThermoFisher Scientific, Cat#34096, Waltham, MA, USA) and exposed to X-ray films (Cat#BL1-810-100) or iBright Imaging System (ThermoFisher Scientific, Waltham, MA, USA). Films were scanned, and band optical densities were quantified using ImageJ software (v1.54j).

### 2.3. Immunofluorescence imaging

Immunofluorescence staining for bromodeoxyuridine (BrdU) was performed to assess cell proliferation, while cleaved caspase-3 (CC3) staining was used to evaluate apoptosis. Cells were treated with the indicated drugs for 48 hours - 72 hours prior to staining. For BrdU immunofluorescence, cells were incubated with 10 nM BrdU (Cat#B23151, ThermoFisher Scientific, Waltham, MA, USA) for 4 h and harvested. The harvested cells were fixed in Cytospin™ Collection Fluid (Cat# 6768001, ThermoFisher Scientific), and cytospun onto positively charged glass slides, permeabilized with 0.1% Triton X-100 in PBS, and subjected to DNA denaturation with 1 N HCl. After blocking with 5% normal goat serum, cells were incubated with anti-BrdU (1:500, Cat#5292S, Cell Signaling Technologies) or anti-CC3 (1:500, Cat#9661S, Cell Signaling Technologies) primary antibody overnight at 4 °C, followed by incubation with a Cy3-labeled secondary antibody. Nuclei were counterstained with DAPI (Roche Life Sciences, Sigma-Aldrich, Cat#10236276001, MI, USA), and slides were mounted using ProLong Gold Antifade Mountant (ThermoFisher Scientific, Cat#P36930). Images were acquired using a Nikon Motic AE31 microscope with NIS Elements software (version F5.21.00). Image analysis was performed in a blind manner. Both the number of immune-positive cells and the total number of cells (defined as DAPI-positive nuclei) were quantified using ImageJ.

### 2.4. Cell growth curve and counting

Cells were seeded in 24-well plates at equal densities (five replicate wells per group) on day 0. At the indicated time points, cells from one well of each replicate were harvested and counted using the Muse™ Cell Analyzer (Merck Millipore) with the Guava® ViaCount™ Reagent (Cat#4000-0041, Cytek Biosciences) according to the manufacturer’s instructions. Cell numbers were recorded daily for 4–5 consecutive days depending on the proliferation rate of each cell line.

Growth curves were generated by plotting the mean cell number ± SD for each time point. Statistical significance between groups was determined using a two-tailed Student’s T-test based on the cell counts obtained on the final day of measurement. A P value < 0.05 was considered statistically significant.

### 2.5. Intracranial xenograft tumors and In Vivo Imaging

All animal procedures were conducted in accordance with the Guide for the Care and Use of Laboratory Animals (NIH Publication No. 86-23, revised 1985) and were approved by the Johns Hopkins University Institutional Animal Care and Use Committee (IACUC), in compliance with the U.S. Animal Welfare Act and Public Health Service Policy.

Intracranial orthotopic xenograft models were established using the luciferase incorporated ATRT cell lines MAF737 and CHLA06. Briefly, mice were anesthetized, and tumor cells were implanted into the brain as previously described^18^.

Mirdametinib was administered orally at 15 mg/kg in 0.5% hydroxypropyl methylcellulose and 0.2% Tween 80, consistent with prior mouse in vivo studies^11,19^. For mirdametinib single-agent studies, mice were treated five times per week. For combination studies, mirdametinib was administered three times a week. Abemaciclib was administered at 50 mg/kg and dissolved in 1% hydroxyethyl cellulose (HEC) in phosphate buffer (pH 2.5). The drug was delivered by oral gavage five times per week. Treatment was initiated 10 days post-implantation for MAF737-Luc xenografts and 3 days post-implantation for CHLA06-Luc xenografts, after confirming tumor establishment by bioluminescence imaging.

Tumor growth was monitored using bioluminescence imaging. Prior to imaging, mice were injected intraperitoneally with 250 μL D-luciferin and anesthetized.

Imaging was performed using an IVIS Spectrum Imaging System (PerkinElmer) with a field of view (FOV) of 22.2 cm and f/1 aperture.

Images were acquired using an Andor iKon camera (IS1651N7095) and analyzed with Living Image software (version 4.7.3, PerkinElmer). Exposure times were adjusted according to signal intensity (3 s for CHLA06 and 120 s for MAF737 xenografts). Total photon flux (photons/sec) within a defined region of interest (ROI) was used for quantitative analysis.

## 3. Results

### 3.1 Mirdametinib inhibits the MAPK pathway and suppresses proliferation in ATRT

We tested mirdametinib against a panel of ATRT representing the three major subgroups, MYC, TYR (tyrosinase), and SHH (sonic hedgehog)^20^. Mirdametinib at nanomolar concentrations suppressed the MAP kinase pathway in all cell lines tested as measured by pERK western blot (Figure 1A).

**Figure 1.**
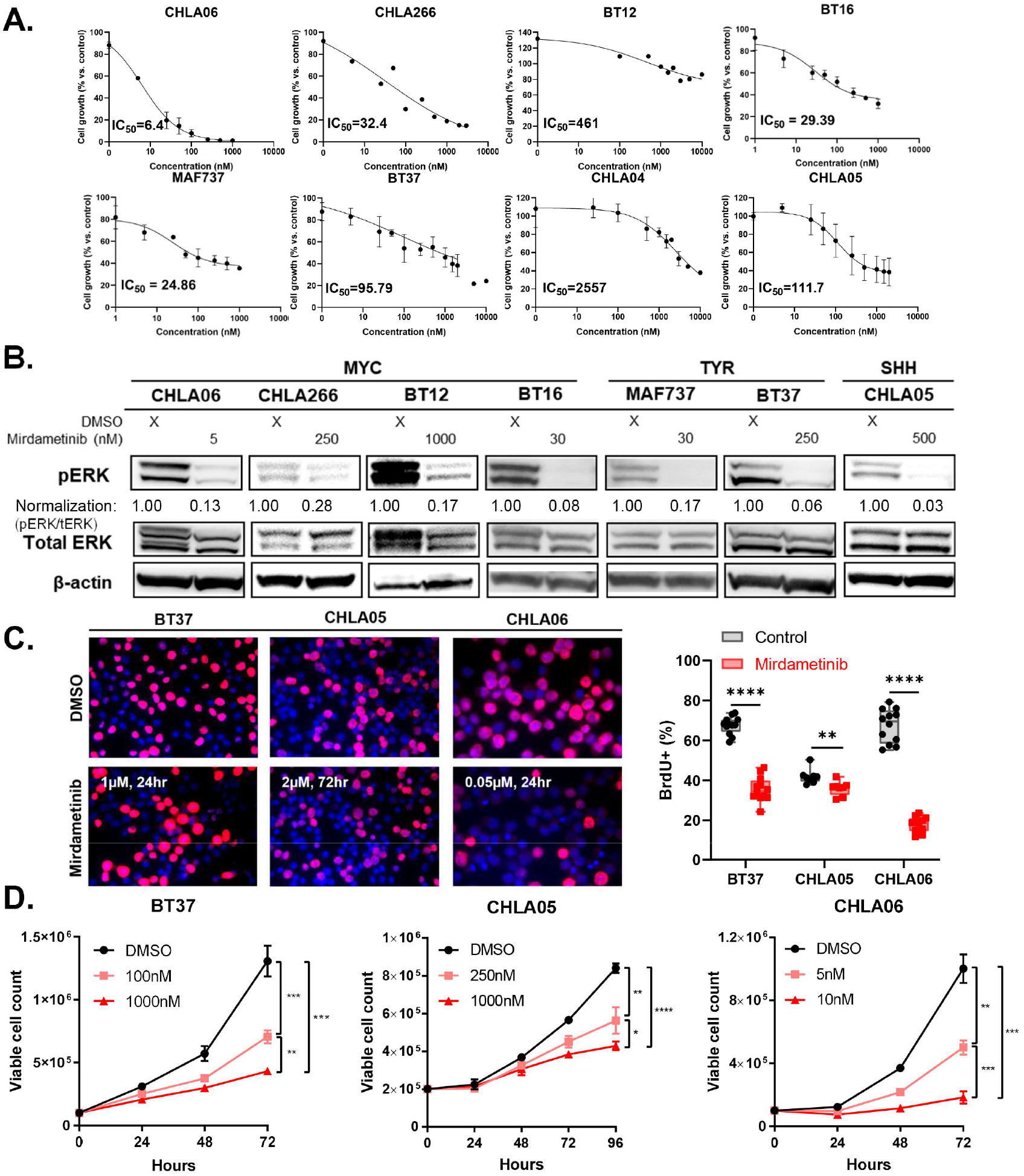
Mirdametinib inhibits the MAPK pathway and suppresses proliferation in ATRT. A. Dose–response curves showing the effect of mirdametinib on cell viability in ATRT cell lines. IC_50_ values were calculated using nonlinear regression model in GraphPad Prism. B. Western blot showing a decrease in the expression of phospho-ERK in ATRT cell lines after treatment with mirdametinib for 24hr-48hr compared to DMSO control. C. Suppression of proliferation as measured by BrdU incorporation for mirdametinib-treated cells and DMSO control. At left are representative immunofluorescence images showing decreased BrdU incorporation. At right is a graph showing the quantification of percent BrdU positivity. The treatment dosing and length are overlayed on the images. **p ≤ 0.01, ****p ≤0.0001, t-test. D. Viable cell count measured by the MUSE Count & Viability assay following mirdametinib treatment.

We determined the IC_50_ for mirdametinib in our ATRT cell lines, and found an IC^50^ range of 10 -2600 nM, though most cell lines were in the low nanomolar range (Figure 1B CHLA06 IC_50_ = 6.4 nM, CHLA266 IC_50_ = 32.4 nM, BT12 IC_50_ = 461 nM, BT16 IC_50_ = 29.4 nM, MAF737 IC_50_ = 24.9 nM, BT37 IC_50_ = 95.8 nM, CHLA04 IC_50_ = 2557 nM, CHLA06 IC_50_ = 111.7 nM).

Mirdametinib suppression of ATRT proliferation was measured by bromodeoxyuridine (BrdU) incorporation (Figure 1C). Mirdametinib decreased BT37 (TYR subtype) BrdU positivity after treatment for 48 hours from 68 percent to 32 percent (p<0.0001 by Student’s t-test). Mirdametinib decreased CHLA05 (SHH subgroup) by 42 to 36 percent(p<0.01). Mirdametinib decreased CHLA06 (MYC subgroup) from 67 to 18 percent (p<0.0001).

Mirdametinib suppressed ATRT growth in a dose-dependent fashion over time as measured by flow cytometric cell viability assay (Figure 1D) in BT37, CHLA05, and CHLA06 (p<0.001 at 72 to 96 hours).

### 3.2 Mirdametinib induces apoptosis in ATRT

Mirdametinib treatment of ATRT for 24 to 72 hours led to an increase in apoptosis as measured by cleaved PARP western blot (Figure 2A). We further measured apoptosis by Annexin V flow cytometry and found that mirdametinib increased the early and late apoptotic fraction significantly compared to controls (BT37 p<0.001; CHLA266 p<0.01 (Figure 2B).

**Figure 2.**
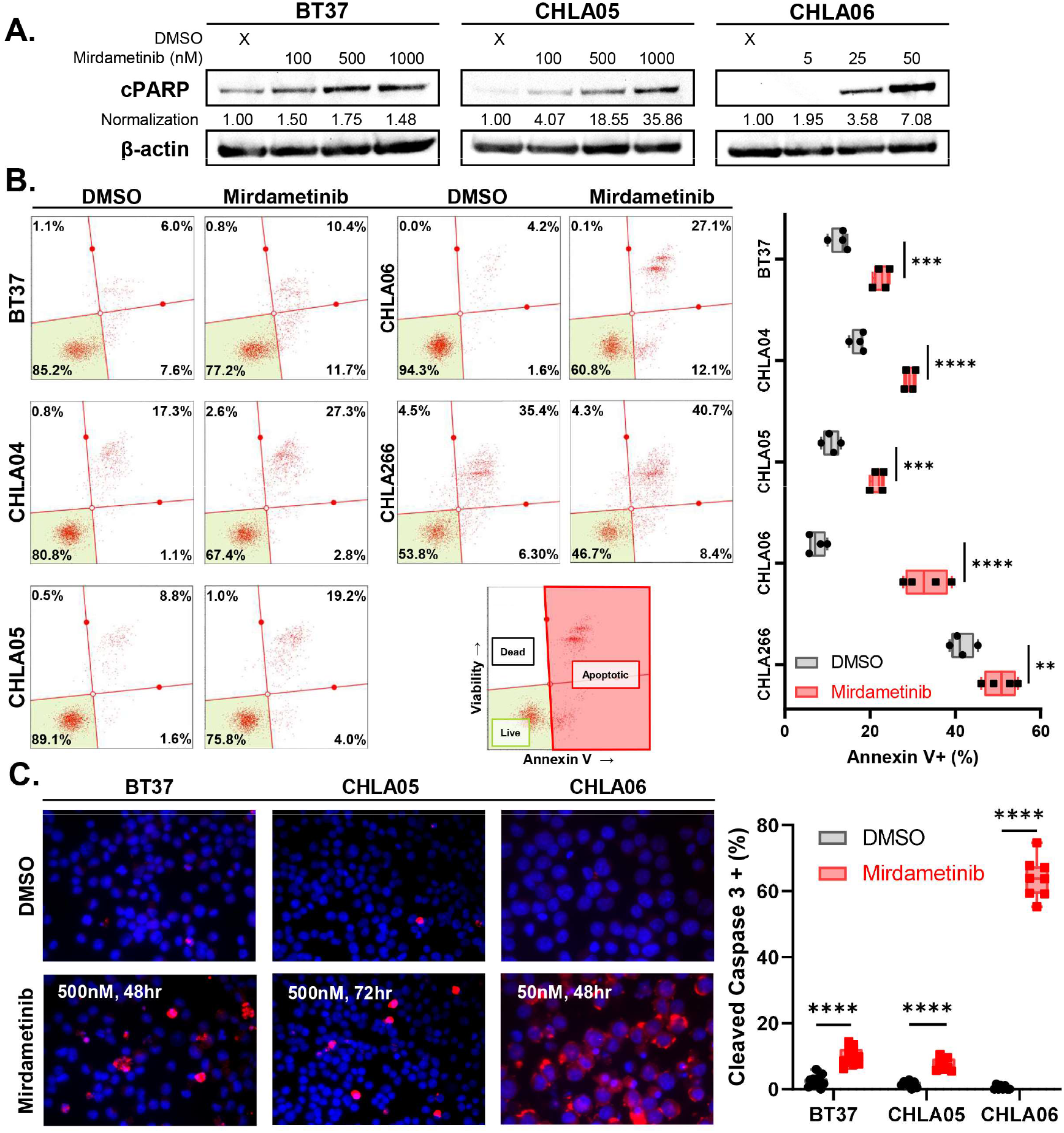
Mirdametinib induces apoptosis in ATRT. A. Western blot showing dose-dependent increase in the expression of cleaved PARP in ATRT cell lines after treatment with mirdametinib for 24hr compared to DMSO control. B. Percent of cells undergoing apoptosis as determined by the MUSE Annexin V assay. The treatment dosing and length for each cell line was: BT37 – 0.5µM for 24hr; CHLA04 - 5µM for 72hr; CHLA05 –0.5µM for 48hr; CHLA06 – 0.05µM for 24hr; CHLA266 – 0.1µM for 48hr. **p ≤ 0.01, ***p ≤0.001, ****p ≤0.0001, t-test. C. Induction of apoptosis as measured by cleaved caspase 3 (CC3) staining for mirdametinib-treated cells and DMSO control. At left are representative immunofluorescence images showing increased CC3 staining. At right is a graph showing the quantification of percent CC3 positivity. The treatment dosing and length are overlayed on the images. ****p ≤0.0001, t-test.

We verified the cytotoxic effect of mirdametinib by performing cleaved caspase 3 immunofluorescence and found statistically significant increases in CC3 positivity after mirdametinib treatment in BT37 (TYR p<0.0001), CHLA05 (SHH p<0.0001) and CHLA06 (MYC p<0.0001) (Figure 2C).

### 3.3 Mirdametinib suppresses MAP kinase pathway in vivo and inhibits tumor growth of ATRT orthotopic xenografts

We then tested mirdametinib against two ATRT orthotopic xenografts. In the TYR subgroup MAF737, 15 mg/kg mirdametinib administered orally 5 days per week delayed tumor growth as measured by bioluminescence at day 23 and day 30 after tumor initiation (Figure 3A p<0.05).

**Figure 3.**
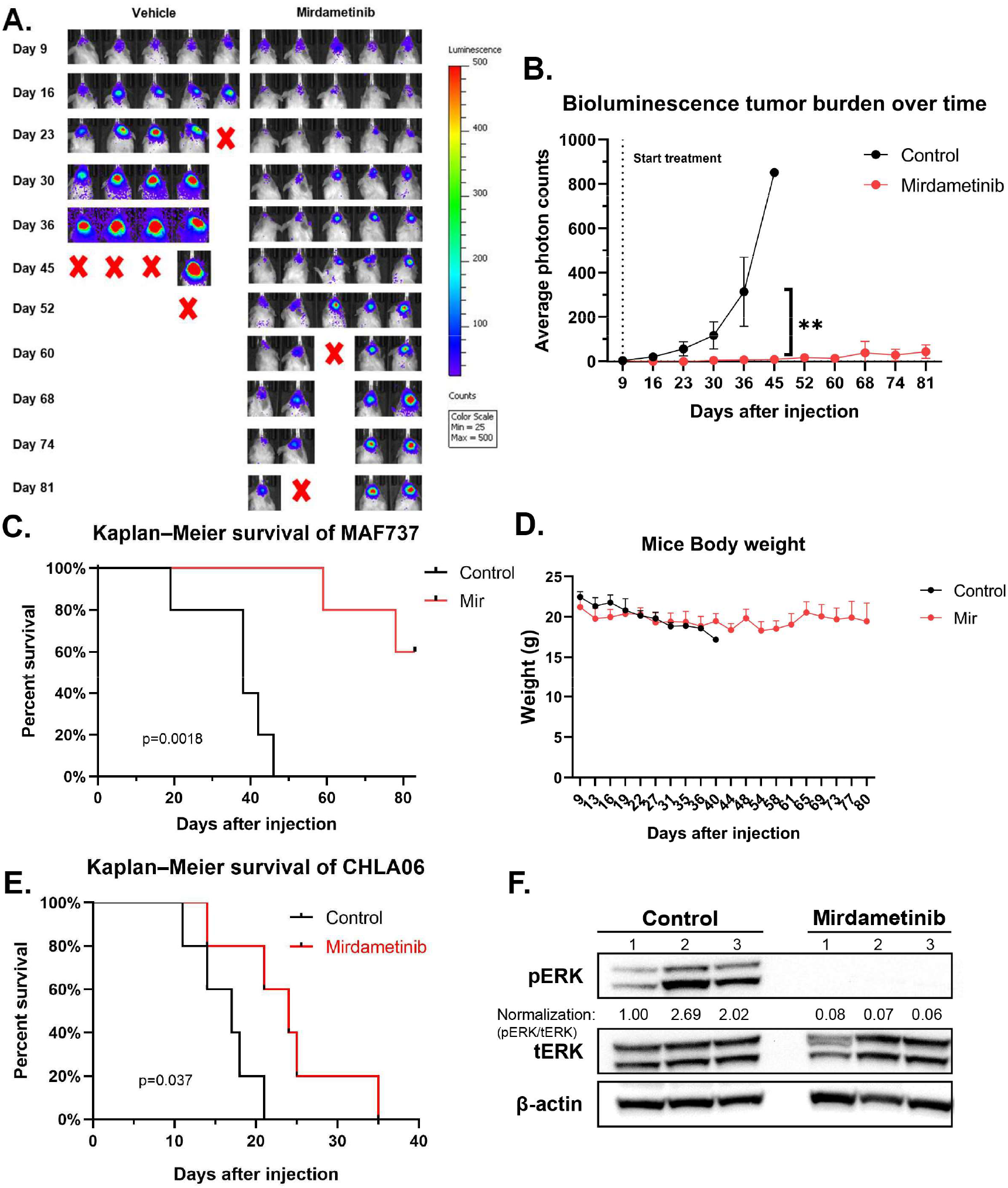
Mirdametinib inhibits the MAPK pathway and extends survival in mice bearing orthotopic MAF737 and CHLA06 tumors. A. IVIS bioluminescence images of control and mirdametinib-treated mice for MAF737. Imaging was performed using identical acquisition settings (binning 8, exposure 120 s, FOV 22.2 cm, f/stop 1). Luminescence intensity is shown in counts (display range 25–500). B. Quantification of bioluminescence tumor burden over time in control and mirdametinib-treated mice. Tumor signal intensity was measured as average photon counts within a defined region of interest using identical imaging settings. C. Kaplan-Meier curve showing survival after injection of MAF737 cells into the deep gray matter of immunodeficient mice. The median survival of control mice (n=5) is 38 days, whereas the median survival of mice treated with mirdametinib (n = 5) was not reached during the observation period(83 days) (p=0.0018 by log rank test). D. Body weight changes over time in control and mirdametinib-treated mice. Body weight measurements obtained within 5 days prior to euthanasia were excluded from analysis. E. Kaplan-Meier curve showing survival after injection of CHLA06 cells into the deep gray matter of immunodeficient mice. The median survival of mice (n=5 in each group) treated with mirdametinib was 24 days compared to 17 days for control mice (p=0.037 by log rank test). F. Western blot showing decreased expression of phospho-ERK in orthotopic CHLA06 tumors compared to control.

Mirdametinib improved overall survival significantly without inflicting toxicity-induced weight changes. Log-rank test showed a significance of p=0.0018 with a more than doubling of median survival (control 38 vs mirdametinib median survival not reached at 75 days) (Figure 3C). Mice were weighed at least weekly, and there was no significant difference in mouse weight between control and treated mice at day 17 of treatment (p=0.4889 Figure 3D).

We extended these results to the highly aggressive MYC subtype ATRT CHLA06 and found that mirdametinib treatment extended survival (p=0.037 by Log-rank test). Median survival improved from 17 days in control to 24 days in mirdametinib treated groups (Figure 3E). At the humane endpoints of the experiment, we treated mice with mirdametinib or control and removed the tumor area of the brain (right cortex and striatum). Western blot analysis demonstrated that mirdametinib treatment reduced MAP kinase pathway activity as measured by phospho-ERK, indicating on-target pharmacodynamics of mirdametinib in ATRT (Figure 3F).

### 3.4 Mirdametinib and the CDK 4/6 inhibitor abemaciclib combine to suppress ATRT growth

While the single agent efficacy of mirdametinib in ATRT orthotopic xenografts reached statistical significance, we sought to investigate if combination therapy would further suppress ATRT growth with non-overlapping toxicity. The significant downregulation of proliferation as measured by BrdU incorporation led us to hypothesize that abemaciclib, a brain-penetrant cyclin dependent kinase 4/6 (CDK4/6) inhibitor currently in clinical trials for pediatric high grade glioma, would combine with mirdametinib to suppress ATRT growth.

Abemaciclib reduces ATRT growth as a single agent, with IC_50_ ranging from 5 nM to 70 nM (Figure 4A). Mirdametinib and abemaciclib combination treatment suppressed phosphor-Rb, a measure of S-phase entry, in CHLA06, BT16 (MYC), MAF737 and BT37(TYR), as shown by Western blot (Figure 4B). Similarly, abemaciclib and mirdametinib combined to reduce BrdU incorporation in BT37 (p<0.05 combination vs control) and CHLA06 (p<0.01 combination vs control) (Figure 4C). We additionally measured BrdU incorporation by flow cytometry in MAF737 (p<0.001) and BT16 (p<0.01), and we found similar reduction in BrdU incorporation with combination therapy compared to control (Figure 4D).

**Figure 4.**
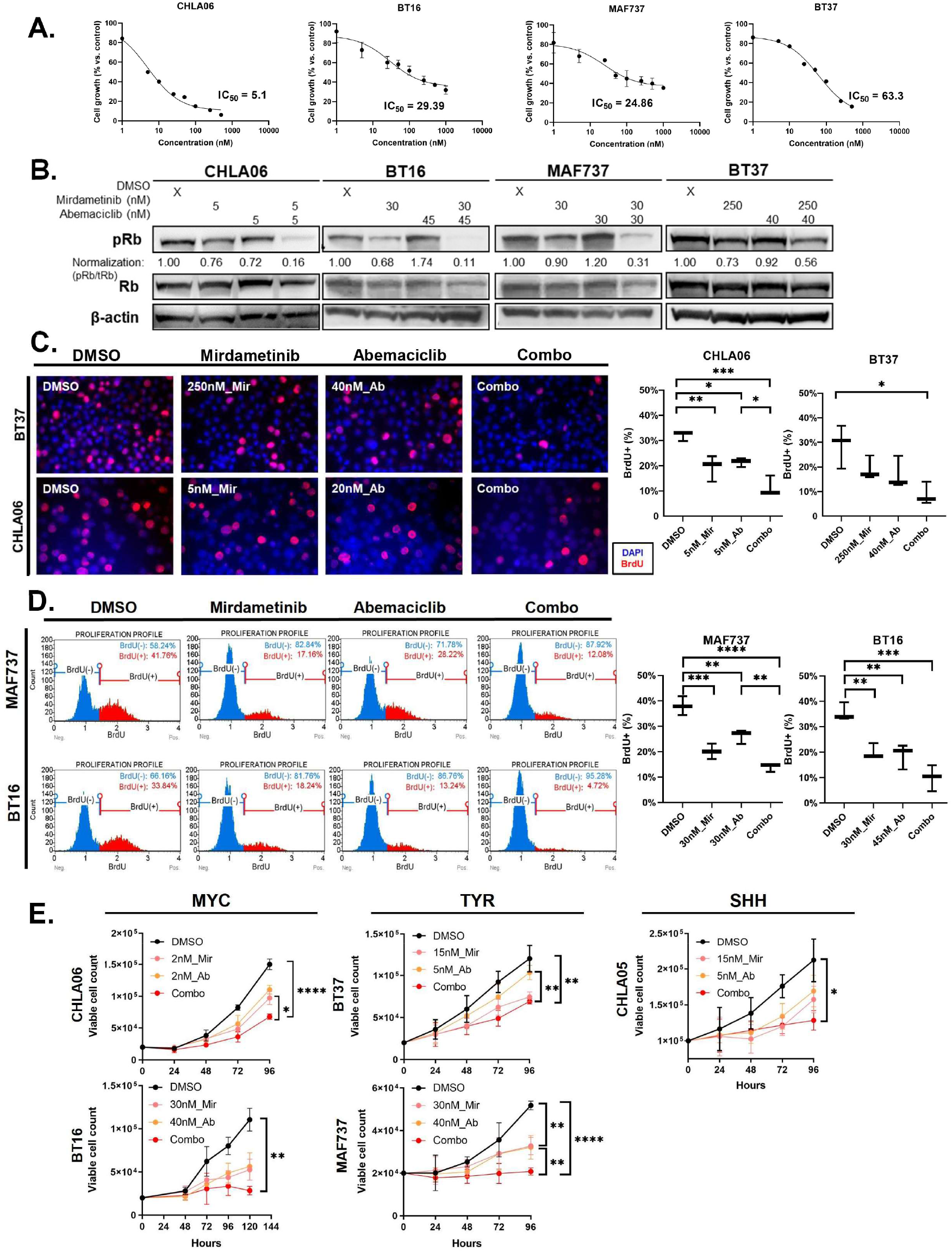
Mirdametinib and CDK 4/6 inhibitor abemaciclib exhibit combinatorial proliferation inhibition in ATRT. A. Dose–response curves showing the effect of abemaciclib on cell viability in ATRT cell lines. IC_50_ values were calculated using nonlinear regression model in GraphPad Prism. B. Western blot showing a decrease in the expression of phospho-Rb in ATRT cell lines after combination treatment with mirdametinib and abemaciclib after 24-48hr. C. Suppression of proliferation as measured by BrdU immunofluorescence incorporation for mirdametinib-treated cells and DMSO control. On the left there are representative immunofluorescence images showing decreased BrdU incorporation. On the right is a graph showing the quantification of percent BrdU positivity by immunofluorescence and D. Suppression of proliferation as measured by BrdU flow cytometry. Quantification of BrdU incorporation by flow cytometry. The treatment dosing and length are overlayed on the images. *p ≤ 0.05, **p ≤ 0.01, ***p ≤ 0.001, ****p ≤0.0001, one-way ANOVA. E. Viable cell count measured by the MUSE Count & Viability assay following single-agent treatment with mirdametinib / abemaciclib or double-agent treatment.

Mirdametinib and abemaciclib combined to reduce ATRT growth compared to control and monotherapy in CHLA06 (p<0.0001), BT16 (p<0.01), MAF737 (p<0.0001), BT37 (p<0.01) and CHLA05 (p<0.05) as determined by flow cytometry viability measurements (Figure 4E).

### 3.5 Mirdametinib and abemaciclib inhibit ATRT tumor growth in orthotopic xenografts with minimal toxicity

We tested mirdametinib and abemaciclib combination against CHLA06 orthotopic xenografts and found that combination therapy significantly decreased tumor progression as measured by bioluminescence at day 3 after initiation (p<0.05).

Overall survival improved with combination therapy compared to monotherapy or control treatment, with median survival increasing from 17 days for control to 33 days for combination therapy (Figure 5B). Of note, we reduced the frequency of treatment with mirdametinib from 5 days per week in the initial experiment in Figure 3E to 3 days per week to improve drug tolerability. We did not detect a difference over the first 30 days of the treatment time course in the weights of mice treated with combination therapy compared to control (Figure 5C). Western blot analysis of tumor collected 4 hour after treatment demonstrated that mirdametinib treatment reduced pERK compared with control tumors (Figure 5D).

**Figure 5.**
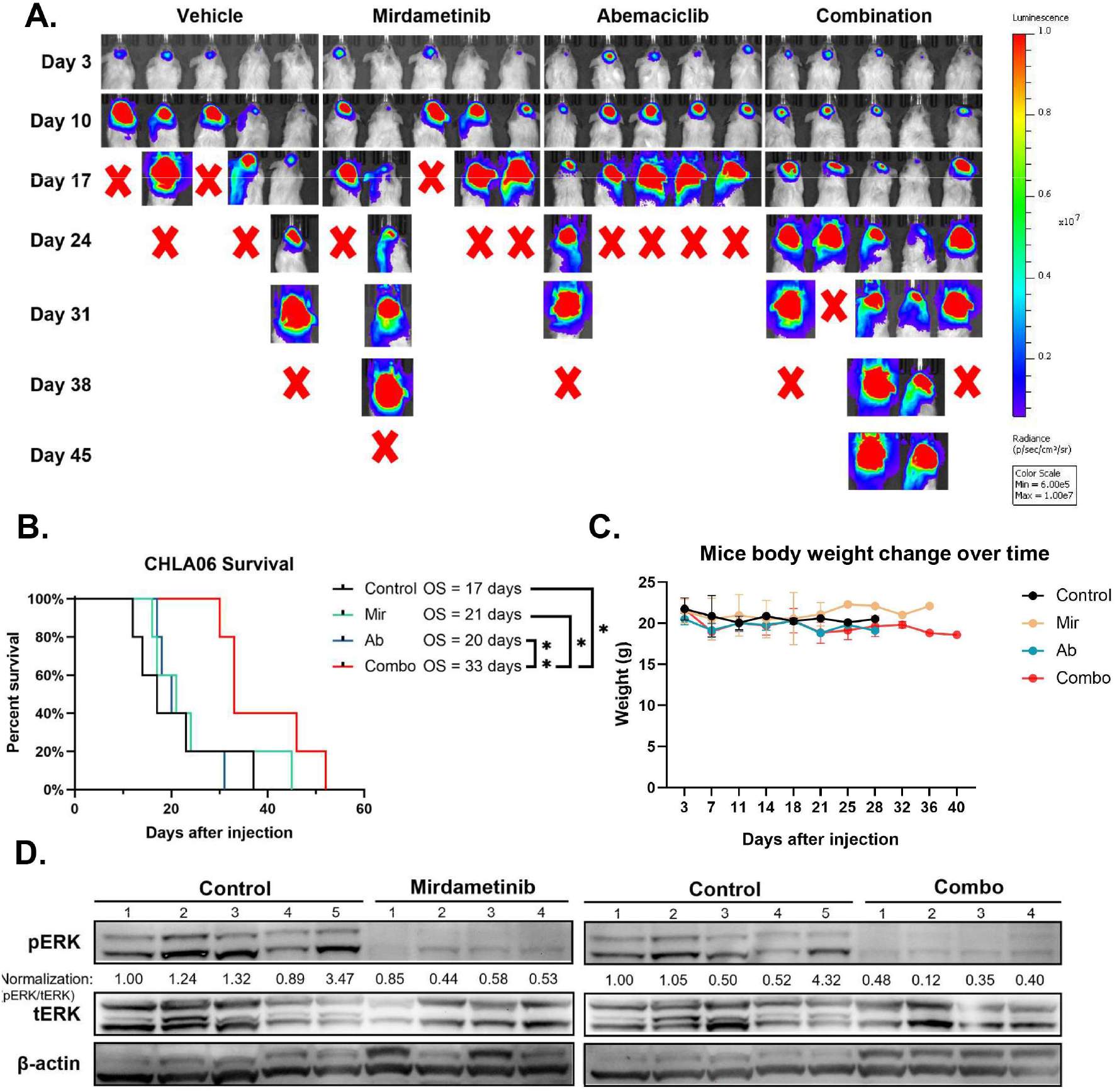
Mirdametinib and abemaciclib extends survival in mice bearing orthotopic CHLA06 tumors. A. IVIS bioluminescence images of control and mirdametinib-treated mice for CHLA06. Imaging was performed using identical acquisition settings (binning 4, exposure 3 s, FOV 22.2 cm, f/stop 1). Luminescence intensity is shown in counts (display range 6e5–1e7). B. Kaplan-Meier curve showing survival after injection of CHLA06 cells into the deep gray matter of immunodeficient mice. The median survival of control mice (n = 5) is 17 days, whereas the median survival of mice treated with mirdametinib and abemaciclib (n = 5) was not reached during the observation period(33 days) *p ≤ 0.05, **p ≤0.01, log rank test. C. Body weight changes over time in control and mirdametinib-treated mice. Body weight measurements obtained within 5 days prior to euthanasia were excluded from analysis. D. Western blot showing decreased expression of phospho-ERK in orthotopic CHLA06 tumors compared to control.

## 4. Discussion

Targeting the LIN28 pathway has long been a focus in our laboratory. We initially showed that disrupting LIN28A itself suppressed ATRT growth and tumorigenicity^5^. We subsequently targeted LIN28A and LIN28B’s downstream effectors, and we demonstrated that suppression of HMGA2 decreased ATRT growth and tumorigenicity^15^. However, neither LIN28A or HMGA2 have clinic-ready therapeutics, prompting us to change our strategy to suppressing LIN28A and LIN28B known effectors in the mTOR and MAP kinase pathways. Previously, we showed that the TORC1/2 inhibitor TAK228 also decreased ATRT growth and tumor formation and synergized with platinum-based chemotherapy_21_. We also targeted the MAP kinase pathway with the MEK inhibitors selumetinib and binimetinib_9_. MEK inhibitors not only suppressed proliferation in ATRT but also induced apoptosis. With the advent of mirdametinib, a MEK inhibitor with improved brain penetration and a pediatric formulation_11_, we hypothesized that mirdametinib would penetrate the blood-brain barrier, suppress the MAP kinase pathway and extend survival of mice bearing ATRT orthotopic xenografts.

Mirdametinib is currently in clinical trials for pediatric brain tumors (ClinicalTrials.gov Identifier: NCT04923126). Mirdametinib is FDA approved for treatment of plexiform neurofibromas in children and adults^12^. Mirdametinib is generally well tolerated in children, with a side effect profile consistent with other MEK inhibitors, primarily dermatologic toxicity ^22^. We found that mirdametinib given 5 days per week orally was well tolerated and extended the life of mice bearing orthotopic xenografts. Mirdametinib effectively suppressed the MAP kinase pathway in the orthotopic xenografts as measured by phospho-ERK western blot.

Prior studies have demonstrated efficacy of CDK4/6 inhibitors in ATRT, particularly in association with radiation therapy^23^. Abemaciclib is a novel CDK4/6 inhibitor with FDA approval for high-risk breast cancer ^24^. Importantly, abemaciclib has activity against breast cancer brain metastases, suggesting improved brain penetration compared to other CDK4/6 inhibitors^25^. Abemaciclib is currently under study in a pediatric phase 2 clinical trial for initial chemotherapy for high-grade glioma (ClinicalTrials.gov Identifier: NCT06413706). Based on the strong inhibition of proliferation we observed with mirdametinib, and the lack of RB mutation found in ATRT, we hypothesized that abemaciclib and mirdametinib would combine to suppress ATRT growth.

In assays of proliferation, mirdametinib and abemaciclib combined to reduce phospho-Rb as well as BrdU incorporation, which are measures of S-phase entry. Similarly, mirdametinib and abemaciclib combined to reduce tumor growth over time. Most significantly, the two drugs improved survival of mice bearing ATRT orthotopic xenografts.

These data support the MAP kinase pathway as a vulnerability in ATRT and suggest that mirdametinib could be a useful agent that could be deployed to improve outcomes in ATRT. Combination therapy with abemaciclib was overall well tolerated and improved survival in an orthotopic model. Our findings support further exploration of this combination in ATRT.

